# Emergence of plasmid-borne *erm*(55)-associated macrolide resistance in *Mycobacterium chelonae* and other rapidly-growing non-tuberculous mycobacteria in Europe

**DOI:** 10.64898/2026.07.08.737060

**Authors:** Camille Allam, Yassine Charmat, Salim Agsous, Zeina Awad, Théo Fouchet, Lilou Goncalves, Nour Ben Salem, Corentin Poignon, Faiza Mougari, Nicolas Véziris, Emmanuelle Cambau

## Abstract

Macrolides are key agents for treating infections caused by non-tuberculous mycobacteria (NTM). Nevertheless, chromosomal *erm* genes conferring inducible macrolide resistance are described in some NTM species, such as *Mycobacterium abscessus* and *M. fortuitum*, whereas *M. chelonae* had long been considered as lacking a functional e*rm*. Recent descriptions from the USA and Japan of a new plasmid-borne *erm*(55) (*erm*(55)^P^) in *M. chelonae* and other rapidly growing mycobacteria (RGM) have challenged this assumption. We investigated *erm*(55)^P^ occurrence in clinical RGM referred to the French National Reference Centre for Mycobacteria between 2012 and 2026 by genome screening and *erm*(55)^P^ specific real-time PCR. Positive isolates underwent long-read whole genome sequencing (GridIon, Oxford Nanopore Technologies). Clarithromycin (CLR) minimum inhibitory concentration (MIC) was determined by broth microdilution (RAPMYCO and FRATMYC, Thermo Fisher) and read up to 14 days.

Five clinical isolates showing inducible CLR resistance (MIC range <0.25-64 mg/L on day 3-4 and 128– >128 mg/L on day 14) were positive for *erm*(55)^P^: one *M. chelonae*, three *M. neoaurum*, and one *M. parafortuitum*. *erm*(55)^P^-positive *M. chelonae* genomes from this and previous descriptions did not cluster together in the phylogenetic analysis of 263 genomes. The assembled plasmids showed high similarity to previously reported *erm*(55)-carrying plasmids, especially within the *erm*(55)^P^ region. The upstream sequence of *erm*(55)^P^ showed a secondary structure compatible with a possible translation attenuation mechanism. These findings document the first report of a plasmid-borne *erm*(55) in Europe in *M. chelonae* and other RGM and raise concern about the emergence of plasmid macrolide resistance in NTM.

## Introduction

Macrolides play a central role in the management of infections caused by non-tuberculous mycobacteria (NTM). They are part of the first-line recommended treatment against *Mycobacterium avium* complex, *M. kansasii*, *M. xenopi*, *M. abscessus* when lacking a functional *erm* gene, and *M. chelonae*, and represent one of the only orally available antimicrobials in NTM therapeutics [1,2]. Chromosomal genes encoding erythromycin ribosome methyltransferases (*erm*), confer inducible resistance to macrolides through 23s rRNA methylation (adenine(2058)-N(6))-methyltransferase). Although *erm*(37) was described in *M. tuberculosis* complex, other *erm* genes were mainly found in rapidly growing mycobacteria (RGM): *erm*(38) in *M. smegmatis*, *erm*(39) in *M. fortuitum*, *erm*(40) in *M. mageritense*, and *erm*(41) in *M. abscessus*. *M. chelonae* was, however, considered fully susceptible to macrolides and lacking a functional *erm* gene [3]. Recently, *erm(55)* was described in six *M. chelonae* clinical isolates displaying an inducible macrolide resistance phenotype, in the USA and in Japan [4,5]. It was also detected in five other RGM species: *M. neoaurum*, *M. iranicum*, *M. bacteremicum*, *M. obuense*, and *M. murale* [5,6]. This new *erm* gene was shown to be carried by a plasmid (*erm*(55)^P^) but was also described as integrated into *M. chelonae* chromosome (*erm*(55)^C^) or into a transposon (*erm*(55)^T^). These two latter variants share 86% and 82% nucleotide (nt) homology with *erm*(55)^P^ sequence, respectively [4]. Furthermore, while *erm*(55)^P^ expression induction may be different from other mycobacterial *erm*, due to its plasmid location, its regulatory mechanism has not yet been elucidated. Our study aims to describe the first *erm*(55)-carrying *M. chelonae* identified in Europe, to evaluate the occurrence of *erm*(55) mediated clarithromycin (CLR) resistance in other clinical RGM, and to provide insight into *erm*(55)^P^ expression induction.

## Material and Methods

### Study design and ethical considerations

We screened for the presence of *erm*(55) within the *M. chelonae* genomes that were sequenced at the associate laboratory of the French National Reference Centre for Mycobacteria (CNR-MyRMA) between 2012 and 2026 in the context of epidemiological investigations [7,8], and in all *M. chelonae* isolates with CLR MIC ≥ 4 mg/L on day 3–4 and/or day 14. We also screened isolates from other RGM species that had been previously reported to carry *erm*(55): *M. bacteremicum*, *M. grossiae*, *M. iranicum*, *M. neoaurum, M. obuense*, and *M. murale* [4–6], or to closely related taxa within the same phylogenetic group [9], regardless of CLR MIC. Species-level identification was performed using MALDI-TOF MS Sirius (Smart Flex, Bruker Daltonics, Billerica, MA, USA), and/or by *hsp65* Sanger sequencing. Sex, age, underlying pulmonary diseases, and history of macrolide treatment were collected only for the patient from whom the *M. chelonae* carrying *erm*(55)^P^ (Mc1) isolate was obtained. All procedures were performed in accordance with the 1964 Helsinki Declaration and its later amendments [10].

### Clarithromycin resistance assessment

CLR MIC was determined using RAPMYCO® plates (CE-IVDR, Thermo Fisher Scientific, Waltham, Massachusetts, USA). For isolates carrying *erm*(55), CLR MIC was additionally determined using FRATMYC1 custom-designed plate (RUO, Thermo Fisher Scientific) containing CLR concentrations up to 128 mg/L. Briefly, a 1:100 dilution of a 0.5 McFarland suspension in Calcium-adjusted Mueller – Hinton broth was distributed into plates incubated at 30 °C under aerobic conditions. CLR MICs were read visually on days 3–4, 7, 10, and 14 to assess inducible resistance. *M. chelonae* CIP 104535 (ATCC 35752T) was the reference strain control for MIC experiments. Mutations in the *rrl* gene causing CLR resistance were screened by variant calling in WGS data for isolates displaying a CLR MIC ≥4 mg/L on day 3–4.

### erm(55) RT-PCR

Isolates were tested by real-time PCR (RT-PCR) for the presence of *erm*(55)^P^, *erm*(55)^C^, and *erm*(55)^T^. Genomic DNA was extracted from freshly grown colonies using the DNeasy UltraClean kit (Qiagen, Germantown, MD, USA). Primers and probes were designed by adapting a previously published PCR assay to RT-PCR specifications [4] (*erm*(55)^P^: F 5′-CAAACCCTGCTACGGCCTAT-3′, R 5′-GCAAGTTTCCCACGATGGTG-3′, and probe: 5’ 6-FAM - TCG-GAG-CGC-TTG-GAC-CTG-TCG-CGG-GT - BHQ®-1 3’, *erm*(55)^C^: F 5’ TTCCAACTACCCTGTTCGCC 3’, R 5’ GTG-AAT-GCC-GAT-GCT-GGA-AC 3’, and probe: 5’ TAMRA - ACG-GCC-AGC-TTT-GCG-CGG-TCA-TCA-T - BHQ®-2 3’, *erm*(55)^T^: F 5’ AAAGCTGGCCATGGTACGAA3’, R: 5’ CAG-AAT-TTG-ACG-CAG-ACC-GC 3’, and probe 5’ Yakima Yellow® - CCG-GGG-TGT-GAA-CGC-CGA-TGC-TGG-A - BHQ®-1 3’).

RT-PCR was run on a QuantStudio 3 Real-Time PCR System (Thermo Fisher Scientific) with a final volume of 20 µL: 2 µL of DNA, 18 µL of Sensimix™ II (Bioline, London, United Kingdom), primers (300 nM), probes (150 nM), and ROX dye (50 nM) for fluorescence normalisation. The PCR included an initial denaturation at 95 °C for 10 minutes, followed by 45 cycles of denaturation at 95 °C for 20 seconds and extension at 56 °C for 40 seconds. Each run included a control with nuclease-free water, and a negative control consisting of *M. chelonae* ATCC 35752T DNA known to be devoid of *erm*(55)^P^.

### Retrieval and analysis of publicly available genomes, plasmids, and metagenomes

We searched NCBI for publicly available sequencing read archives (SRA) from *M. chelonae*, *M. neoaurum* and *M. parafortuitum*. FastQ sequences, and geographic location (country) were collected from BioProjects <u>PRJDB1758</u>, <u>PRJDB2512</u>, <u>PRJDB14421</u>, <u>PRJDB35518</u>, <u>PRJEB62124</u>, <u>PRJEB75642</u>, <u>PRJEB75747</u>, <u>PRJEB88541</u>, <u>PRJEB9515</u>, <u>PRJNA1019825</u>, <u>PRJNA1169681</u>, <u>PRJNA323571</u>, <u>PRJNA347845</u>, <u>PRJNA574109</u>, <u>PRJNA657124</u>. All published sequences containing the *erm*(55)^P^ sequence: MCHL-2034 (accession number SAMN33426568) [4], PV530475 to PV682406 [6], MCHL-SRL2021-127 (AP043661), and LC872744 to LC872748 [5], *M. iranicum* H39 (BioProject PRJNA317657), and *M. obuense* UC1 (BioProject PRJNA279894) [6] were retrieved for geographical cartography. *erm*(55)^P^ protein sequence was blasted against MGnify (https://www.ebi.ac.uk/metagenomics/) and Integrative Microbial Genomes & Microbiomes (https://img.jgi.doe.gov/cgi-bin/m/main.cgi) metagenomic databases. Sequences were retained if they showed>80% homology over the total protein length, allowing detection of close homologous proteins. Geographic location and isolation source were collected and visualised using mapchart (https://www.mapchart.net/).

### Whole-genome sequencing (WGS)

For short reads, libraries were prepared from 0.2 ng/μL purified DNA using Illumina DNA Prep (Illumina, San Diego, CA, USA) according to the manufacturer’s instructions and sequenced on an Illumina MiSeq platform (2 × 150 bp) targeting a theoretical depth ≥30×. Isolates positive for *erm*(55) by RT-PCR were additionally sequenced using long-read technology. Libraries were prepared from 400 ng DNA using the ligation sequencing gDNA native barcoding kit and run on a GridION (Oxford Nanopore Technology, Oxford, United Kingdom). Extracted DNA and libraries were quantified using a Quantus Fluorometer (Promega, Charbonnières-les-Bains, France), and purity using a Bioanalyzer and the High Sensitivity DNA Kit with High Sensitivity DNA Chips (Agilent Technologies, Santa Clara, USA).

### Bioinformatic analyses of RGM isolates and plasmids

For all retrieved and *de novo* sequenced genomes, an expected species relative abundance>95% was verified using Kraken2 (v2.17.1 using prebuilt Refseq indexes: core_nt v2024-09-04 database from Galaxy, https://usegalaxy.org). Low-quality reads (Phred score <30) were removed using Trimmomatic (v0.39). Short-read assemblies were generated using SPAdes (v3.15.5) and retained if they met predefined sufficient quality criteria: N50> 40,000; <250 contigs; and genome size constraints compatible with *M. chelonae* genome (exclusion if total length <4.5 Mbp or> 6.5 Mbp, QUAST, v5.3.0). For the phylogenetic tree, *M. chelonae* assemblies were annotated using Prokka (v1.14.6), core-genome alignment generated using Roary (v3.13.0), and phylogenetic tree reconstructed using IQ-TREE (v2.4.0) with 1000 ultrafast bootstrap replicates (-bb 1000) and visualised using iTOL. Long-read FastQ files were quality-checked with NanoPlot (v1.42.0) and assembled with Flye (v2.9.6) and polished by Medaka (v2.1.1). Functional annotation was performed using Bakta (v.1.9.4). Visualisation of plasmids and regions encompassing *erm*(55)^P^ was done using Geneious Prime 2026 (v 2026.1). Plasmid core genes were assessed using Panaroo (v1.5.0) with the --clean-mode strict option and visualised using an in-house R script (v4.5.2). pErm55Mc plasmid was also compared with other plasmids identified in this study and published plasmids: PV530475 to PV682406, and LC872744 to LC872748 using online NCBI BLASTn local alignment and default algorithm parameters.

### Genomic exploration of macrolide resistance

*erm*(55)^P^, *erm*(55)^C^, and *erm*(55)^T^ sequences were blasted against all included assemblies (retrieved from SRA and *de novo* sequences) and NCBI core-nt database (BLASTn local alignment). For *rrl* variant calling, reads were aligned to NCBI reference genomes (*M. chelonae* ATCC 35752T, RefSeq accession NZ_CP010946.1, *M. neoaurum* VKM Ac-1815D, CP006936.2, and *M. parafortuitum* JCM 6367, AP022598.1) using BWA-MEM2 (v2.3, short-reads), or minimap2 (v2.30 long reads). Variants were called using FreeBayes (v1.3.6) with ploidy set to 1, focusing on mutations A2058G/C and A2059G/C (*E. coli* numbering) in the *rrl* gene. Only variants with a quality score>30 were retained.

### BLAST of Erm protein sequences and intergenic regions

Erm(55)^P^ amino-acid (aa) sequence (accession number WP_311033296.1) was aligned to those of other Erm proteins described from *M. tuberculosis* H37Rv (Erm37, RefSeq NP_216504.1), *M. smegmatis* ATCC14468 (Erm38, RefSeq WP_063844518.1), *M. fortuitum* (Erm39 accession number QBK17507.1), *M. mageritense* (Erm40, accession number AAS76623.1), *M. abscessus* ATCC19977 (Erm41, RefSeq WP_012296532.1), *Prescottella* (ex *Rhodococcus*) *equi* (Erm46, accession number WP_087587931.1), and also to the *erm* encoded hypothetical protein from *M. koreensis* JCM19956 (formerly *Mycobacterium* koreense, accession number BBY55415.1) that carries an *erm* gene close to *erm*(55)^P^.

*erm*(55)^P^ upstream intergenic region was aligned against *erm*(37) from *M. tuberculosis* H37Rv (RefSeq NC_000962.3), *erm*(38) from *M. smegmatis* strain Jucho (accession number NZ_CP080274.1), *erm*(39) from *M. fortuitum* (accession number NZ_AP025518.1), *erm*(40) from *M. mageritense* AY570506.1 (*pgaE*, partial coding sequence; putative transcriptional regulator and *erm*(40) genes), *erm*(41) from *M. abscessus* ATCC19977 (RefSeq NC_010397.1), intergenic region of *P. equi* strain PAM2287 plasmid pRErm46 (accession number CP095479.1), and upstream *erm* from *M. koreensis* JCM19956 (BioProject <u>PRJDB7717</u>). Alignments were performed using MAFFT online tool (auto mode, https://mafft.cbrc.jp/alignment/server/index.html) generating a Clustal alignment file and visualised in Jalview 2.11.5.1.

### Analysis of plasmid region upstream erm(55)

The annotated protein sequence immediately upstream *erm*(55)^P^ was searched against the UniProt database (https://www.uniprot.org/) to identify homologous annotated proteins and queried in AlphaFold (https://alphafold.ebi.ac.uk/) to assess the protein structure. A functional annotation was performed using eggNOG-mapper v2.1.13 (diamond BLASTp version BLOSUM62, <0.05 retained e-value and 50% query coverage) retrieving orthologs and clusters of orthologous (COG) categories. The intergenic upstream *erm*(55)^P^ region was analysed using ORF finder (NCBI) to find a putative leader peptide. RNA fold (http://rna.tbi.univie.ac.at/cgi-bin/RNAWebSuite/RNAfold.cgi) was used to define the putative secondary structure of mRNA, minimum free energy and thermodynamic ensemble prediction. Putative ribosome binding site (RBS) sequences of *erm*(55) and the leader peptide were predicted from the 17 nucleotide sequence upstream start codons using RBS Strength Prediction & Design Tool (https://novoprolabs.com/tools/rbs/). Then RNA canvas (https://rnacanvas.app/) was used for intergenic structure visualisation.

## Results

### Inducible macrolide resistance in an erm(55)-carrying M. chelonae, and other RGM

From screening of our *M. chelonae* genomes collection, we obtained 1/49 sequence (Mc1) with a positive blast for *erm*(55): 100% nt identity for *erm*(55)^P^, 86.3% for *erm*(55)^C^, and 83.4% for *erm*(55)^T^. *erm*(55)^P^ RT-PCR positivity confirmed this result. Mc1 was isolated in 2020 from the sputum sample of a woman suffering from *M. avium* pulmonary disease, and this was considered as *M. chelonae* colonisation. She had previously received a two-year treatment with azithromycin and ethambutol. A 2-month oral treatment with azithromycin, clofazimine, and amikacin liposomal inhalation suspension was initiated in 2020, after the *M. avium* and *M. chelonae* concomitant isolation. Mc1 was no longer isolated but *M. avium* was isolated in 2024 and displayed an A2058G *rrl* mutation with heteroresistance leading to CLR resistance, and a A1408G *rrs* mutation leading to amikacin resistance. Neither *erm*(55)^P^ nt sequence, nor any sequences of the plasmid *erm*(55) matched *M. avium* consecutive genomes. Analysis of 263 *M. chelonae* genomes (214 retrieved from public SRA, 49 from our in-house collection) showed an absence of clustering between *M. chelonae erm*(55)^P^ carrying isolates (MCHL-2034 from the USA, MCHL-SRL2021-127, from Japan, and MCHL-2632 from France, Figure 1A). Regarding other RGM, 4/18 included isolates tested positive for *erm*(55)^P^ by RT-PCR: three *M. neoaurum* (Mn1, Mn2 and Mn3) and 1 *M. parafortuitum* (Mp1, Table 1). Mn1 and Mn2 were isolated from blood cultures (central catheter source) in two patients, with one being suspected of infectious endocarditis. Mn3 and Mp1 were isolated from respiratory samples and considered as colonisation.

**Figure 1:**
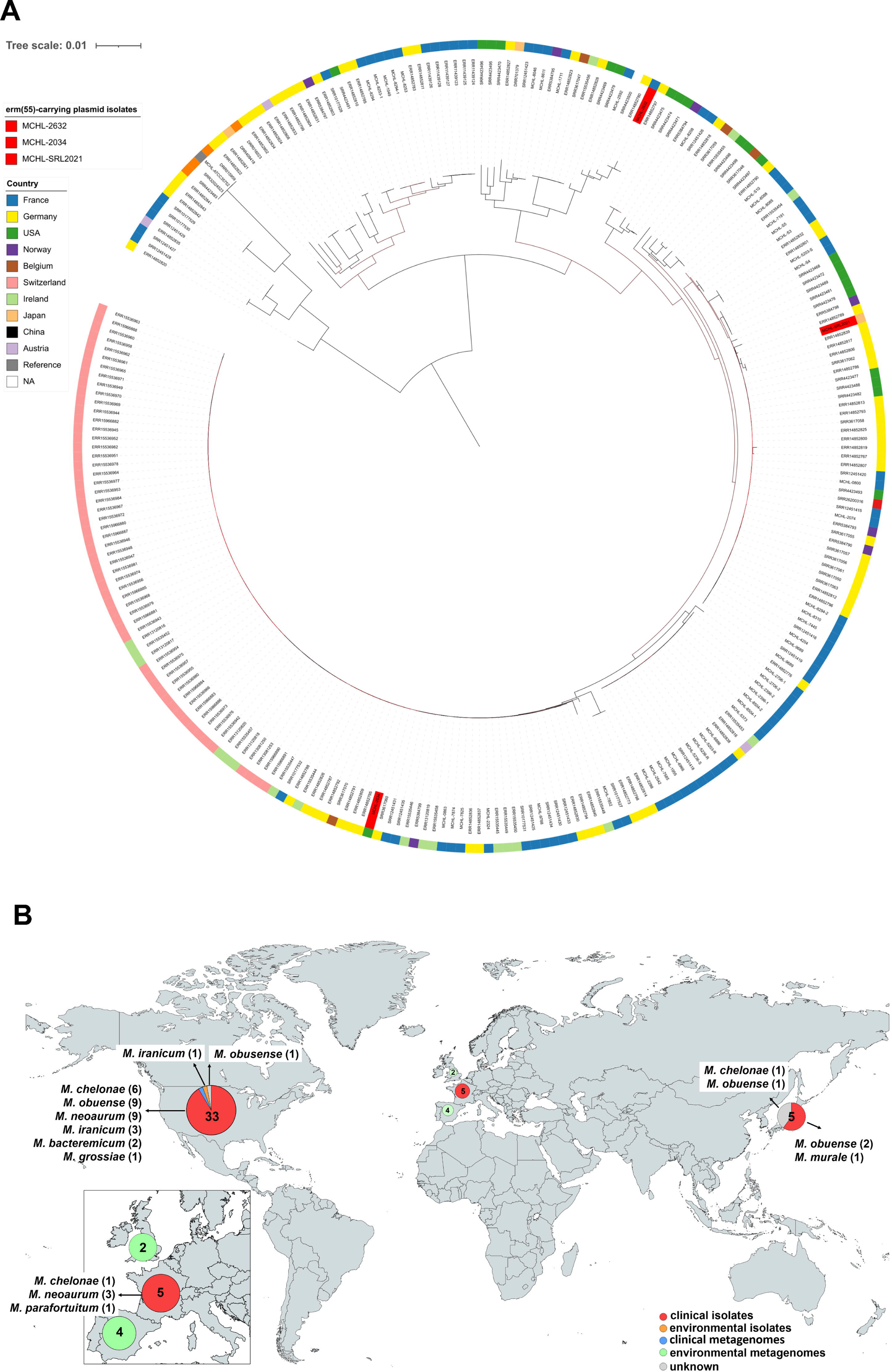
Distribution of *erm*(55)^P^ gene and phylogeny of *erm*(55)^P^ carrying *M. chelonae* isolates. Core-genome alignment phylogenetic tree of retrieved and *de novo* sequenced *M. chelonae* genomes according to their geographical location and *erm*(55)**^P^** carrying status (A). Worldwide geographic distribution of *erm*(55)**^P^** carrying genomes or metagenomes from clinical or environmental sources (B).

**Table 1.**
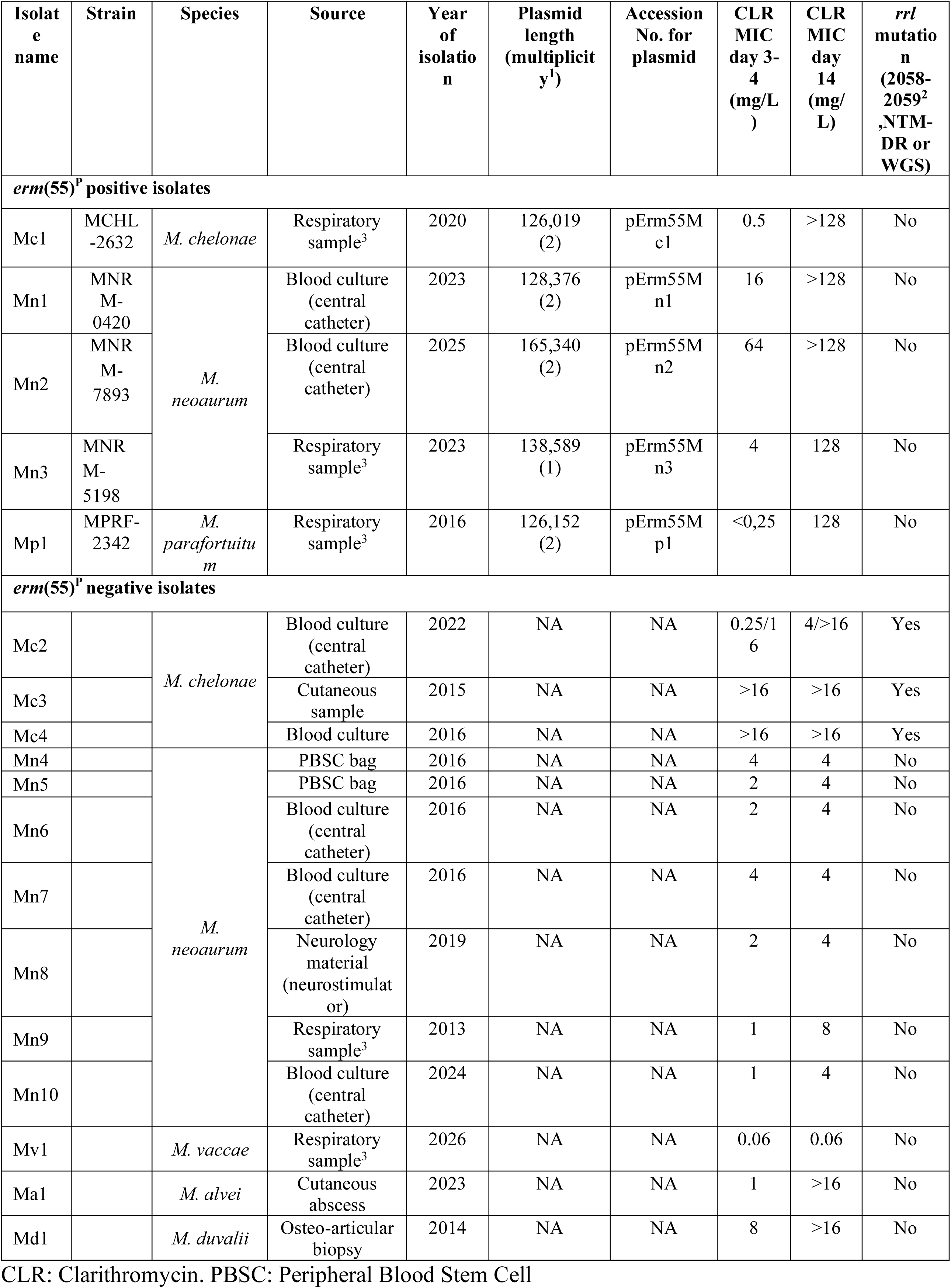
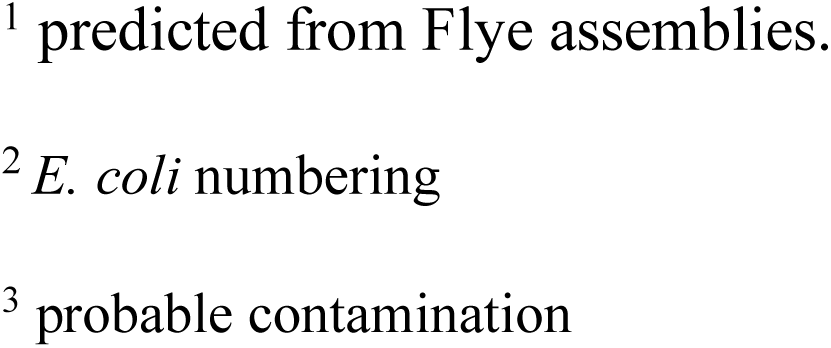
Description of the mycobacterial isolates showing a day 3-4 CLR MIC ≥4 mg/L and/or a day 14 CLR MIC ≥4 mg/L included in this study.

| Isolate name | Strain | Species | Source | Year of isolation | Plasmid length (multiplicity<sup>1</sup>) | Accession No. for plasmid | CLR MIC day 3-4 (mg/L) | CLR MIC day 14 (mg/L) | *rrl* mutation (2058-2059<sup>2</sup>, NTM-DR or WGS) |
| --- | --- | --- | --- | --- | --- | --- | --- | --- | --- |
| ***erm(55)*<sup>P</sup> positive isolates** |  |  |  |  |  |  |  |  |  |
| Mc1 | MCHL-2632 | *M. chelonae* | Respiratory sample<sup>3</sup> | 2020 | 126,019 (2) | pErm55Mc1 | 0.5 | >128 | No |
| Mn1 | MNR M-0420 | *M. neoaurum* | Blood culture (central catheter) | 2023 | 128,376 (2) | pErm55Mn1 | 16 | >128 | No |
| Mn2 | MNR M-7893 |  | Blood culture (central catheter) | 2025 | 165,340 (2) | pErm55Mn2 | 64 | >128 | No |
| Mn3 | MNR M-5198 |  | Respiratory sample<sup>3</sup> | 2023 | 138,589 (1) | pErm55Mn3 | 4 | 128 | No |
| Mp1 | MPRF-2342 | *M. parafortuitum* | Respiratory sample<sup>3</sup> | 2016 | 126,152 (2) | pErm55Mp1 | <0,25 | 128 | No |
| ***erm(55)*<sup>P</sup> negative isolates** |  |  |  |  |  |  |  |  |  |
| Mc2 |  | *M. chelonae* | Blood culture (central catheter) | 2022 | NA | NA | 0.25/16 | 4/>16 | Yes |
| Mc3 |  |  | Cutaneous sample | 2015 | NA | NA | >16 | >16 | Yes |
| Mc4 |  |  | Blood culture | 2016 | NA | NA | >16 | >16 | Yes |
| Mn4 |  | *M. neoaurum* | PBSC bag | 2016 | NA | NA | 4 | 4 | No |
| Mn5 |  |  | PBSC bag | 2016 | NA | NA | 2 | 4 | No |
| Mn6 |  |  | Blood culture (central catheter) | 2016 | NA | NA | 2 | 4 | No |
| Mn7 |  |  | Blood culture (central catheter) | 2016 | NA | NA | 4 | 4 | No |
| Mn8 |  |  | Neurology material (neurostimulator) | 2019 | NA | NA | 2 | 4 | No |
| Mn9 |  |  | Respiratory sample<sup>3</sup> | 2013 | NA | NA | 1 | 8 | No |
| Mn10 |  |  | Blood culture (central catheter) | 2024 | NA | NA | 1 | 4 | No |
| Mv1 |  | *M. vaccae* | Respiratory sample<sup>3</sup> | 2026 | NA | NA | 0.06 | 0.06 | No |
| Ma1 |  | *M. alvei* | Cutaneous abscess | 2023 | NA | NA | 1 | >16 | No |
| Md1 |  | *M. duvalii* | Osteo-articular biopsy | 2014 | NA | NA | 8 | >16 | No |

CLR: Clarithromycin. PBSC: Peripheral Blood Stem Cell
<sup>1</sup> predicted from Flye assemblies.
<sup>2</sup> *E. coli* numbering
<sup>3</sup> probable contamination

### erm(55) **^P^** distribution in publicly available bacterial genomes and metagenomes

We further explored the e*rm*(55)^P^ distribution across NTM genomes. Apart from previously published plasmids from the USA and Japan, *erm*(55)^P^ protein sequence yielded positive BLAST hits in two mycobacterial genomes (*M. iranicum* H39 and *M. obuense* UC1, 100% identity) [6], and for metagenomes from three locations (one skin microbiome from the USA, 100% identity, ERP116514 study, one hospital wastewater from the United Kingdom, ERP123561, 100% identity, and one wastewater from Spain, 81.5% identity, ERP131768, Figure 1B). e*rm*(55)^P^ protein sequence showed 74% aa identities with a genome of a clinical *M. koreensis* (JCM 19956, human sputum, Seoul, Korea). No plasmid was identified from the long-read assembly of this genome. Finally, only 20 WGS SRA of *M. neoaurum* and 8 of *M. parafortuitum* were publicly available; none of them carried *erm*(55)^P^. Database search for protein sequences corresponding to the genes *erm*(55)^C^ *erm*(55)^T^, which show 77% and 82% identity with e*rm*(55)^P^ protein sequence, yielded no additional positive BLAST hits.

### Phenotypic and genotypic characterisation of RGM carrying erm(55) plasmids

The five *erm*(55)^P^ carrying isolates displayed inducible resistance to CLR: day 3-4 MIC range was <0.25-64 mg/L according to species, while MICs reached 128 or>128 mg/L after 14 days of incubation (Table 1). Among *erm*(55)^P^ negative isolates, four other *M. chelonae* isolates displaying day 3-4 CLR resistance tested negative for *erm*(55) RT-PCR but showed A2058 or A2058 *rrl* mutations. Two other isolates (Ma1 and Md1) also showed a CLR MIC increase between day-3 and day-14, but no *erm*(55) variant was identified from their genome analysis nor by RT-PCR screening.

Long-read assembly of the five *erm*(55)^P^ carrying isolates showed contigs of 126,019 to 165,340 bp categorised as circular (Table 1, and example of Mc1 plasmid, Figure 2A). The annotation of these contigs confirmed the presence of an 813 bp *erm* gene showing 100% nt identity with *erm*(55)^P^ sequence. None of these isolates harboured *rrl* mutations. We analysed the shared coding and total nucleotide sequences of these five plasmids with the 35 previously published (Figure 1B). Taken together, Panaroo analysis designed 133/368 (36%) genes as core or soft-core genes (95-100% presence) and 235/368 (64%) as shell or cloud genes (<15-95% presence, Figures 2B and 2 C, Table S1). The BLASTn analysis of pErm55Mc1 showed a nucleotide homology range of 97%-100% with 70%-100% query coverage compared to the 39 other plasmids (Table S2). The core genome included the 38 kb conserved Type IV/Type VII secretion system (T4SS/T7SS) region, as previously shown [6]. Interestingly, all but one 348-bp gene of the pErm55Mc1, annotated as a hypothetical protein, belonged to the core genome.

**Figure 2:**
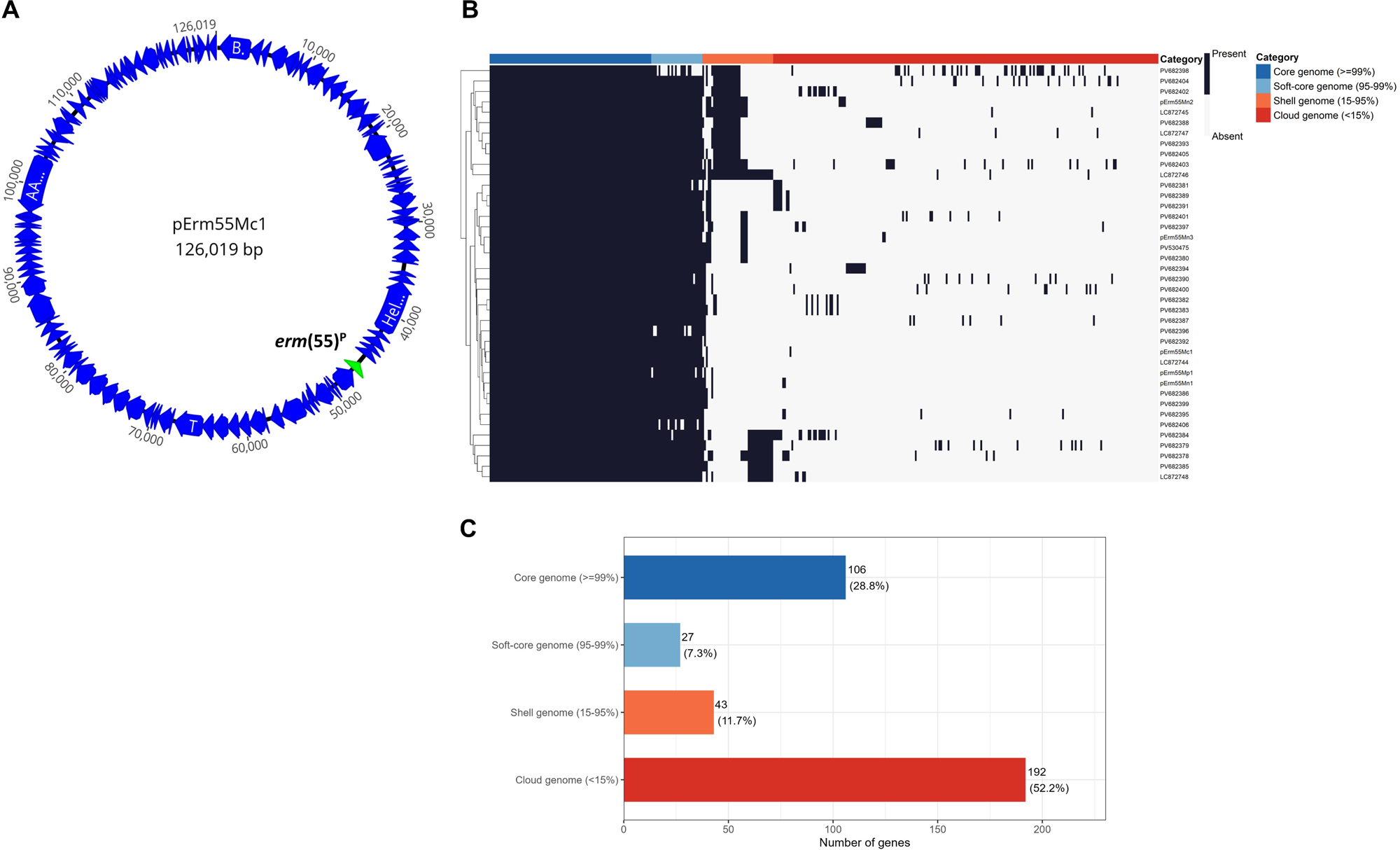
Comparative analysis of the *erm*(55)-carrying plasmids. Details of the pErm55Mc1 *erm*(55)-carrying plasmid gene found in the *M. chelonae* isolate reported in this study (A). The green arrow represents *erm*(55). Hierarchical clustering of the 40 plasmid pangenomes (30 from the USA [8], 5 from Japan [5], and 5 from this study) as identified by Panaroo (B). Plasmid pangenome composition according to Panaroo categories (B).

### Comparison of erm(55) **^P^** to mycobacterial and closely related erm genes

Erm(55)^P^ protein sequence was compared to five mycobacterial 23s methyltransferases encoded by *erm*(37 to 41), to the *M. koreensis* JCM19956 putative methyltransferase (74% aa identity with *erm*(55)^P^ methyltransferase), and to the methyltransferase from *P. equi* encoded by pRErm46 *erm*(46)-carrying plasmids (70% aa homology to *erm*(55)^P^ methyltransferase) [4]. Multiple sequence alignment revealed highly conserved catalytic core regions, particularly in rRNA-interaction domains (aa 21-181), while the other regions and C-terminal regions display greater variability (Figure 3).

**Figure 3:**
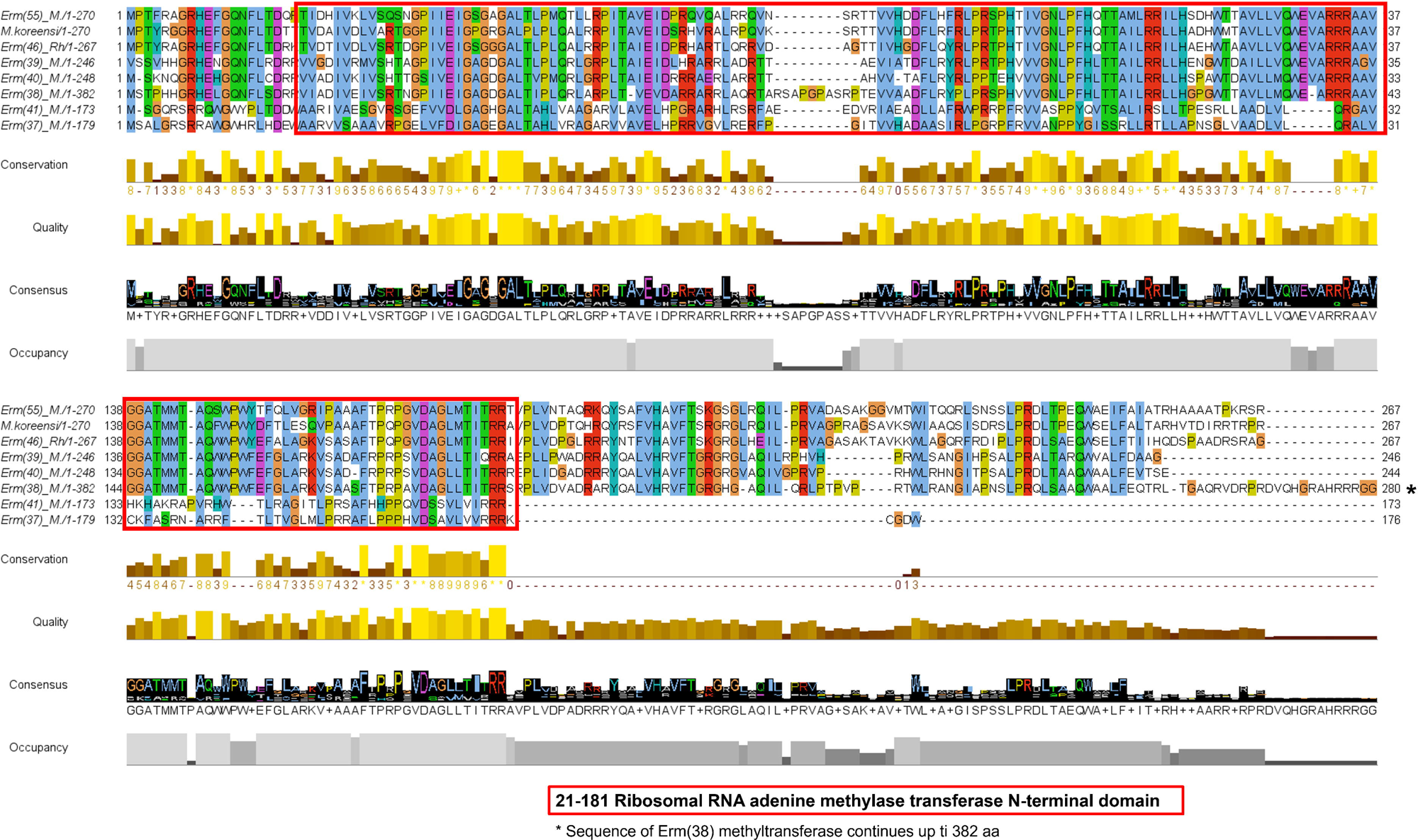
**Alignment of Erm(55)^P^ protein sequence with other mycobacterial Erm proteins and Erm(46) from *P. equi*** MAFFT protein multi-alignment of Erm(55)^P^ first 280 aa to other mycobacterial Erm(37 to 41) and to Erm(46), coloured by clustal colour scheme, and visualised in Jalview. The red box represents the methyltransferase domain, as described in UniprotKB from Erm(37 to 41), and Erm (46). Conservation represents the aa conservation score (out of ten), quality the biochemical proximity between aa, consensus the consensus sequence, and occupancy the proportion of sequences with no gaps at a given position. * The remaining portion of the Erm(38) sequence, from aa 280 to 382, was cut for visualisation purposes

### Exploration of erm(55)**^P^** plasmidic region

We examined the regions encompassing *erm*(55) in the 40 plasmids. The 2,004 bp upstream gene codes for a 668 aa protein with 100% aa identity in 40/40 plasmids (Figure 4A). It was classified as “uncharacterised protein” (UniprotKB), or as “MarR family transcriptional regulator” (AlphaFold). The sequence also matched a *Mycobacterium sp.* SMC-8 plasmid-encoded protein (accession number UXA15786, 97% identity) and a *M. tusciae* hypothetical protein (WP_083129136, 90% homology). These two genomes lack *erm*(55)^P^. EggNOG-mapper functionally annotated the protein as COG1802 (K category, transcriptional regulation, with the closest seed ortholog sharing 50% identity over 97% of the query in M*ycobacterium sp.*). No InterPro domains were identified, suggesting that its function remains uncharacterised despite its presence in other mycobacterial plasmids and genomes.

**Figure 4:**
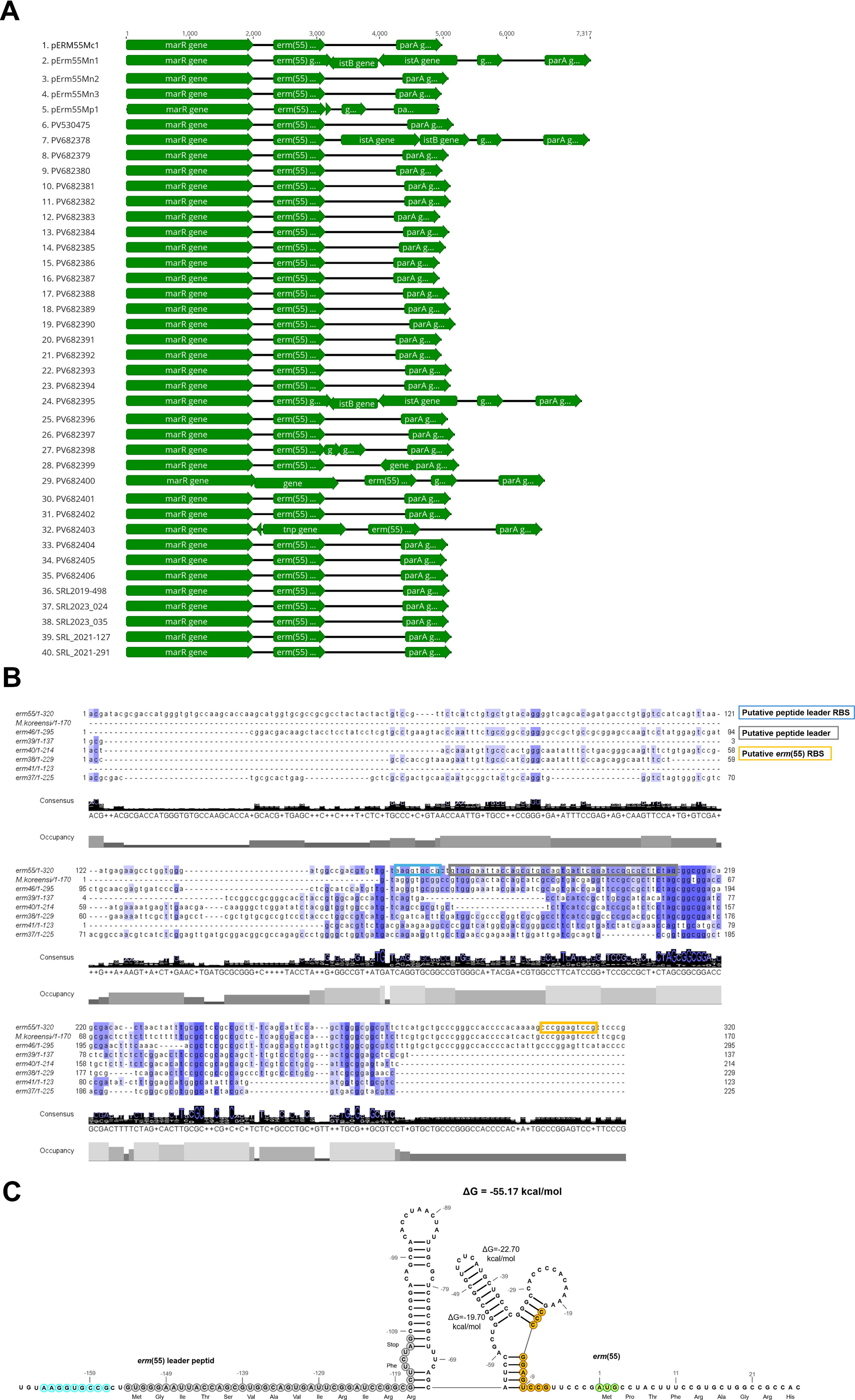
Analysis of intergenic upstream *erm*(55)^P^ plasmid region. Alignment of *marR*-*parA* plasmid regions for the 40 *erm(*55)-carrying plasmids (30 from USA [8], 5 from Japan [5], and 5 from this study) visualised in geneious prime software (A). MAFFT nt alignment of *erm*(55) intergenic upstream regions to other mycobacterial *erm*(37 to 41), putative *M. koreensis erm*, and to *erm*(46), coloured by nt identity (B). The grey box represents the putative leader peptide, along with its putative RBS sequence (blue), and the orange box represents the putative *erm*(55) RBS. Visualisation of *erm*(55) −169 nts upstream mRNA, and RNA fold predicted secondary structure of −117 to −9 nts mRNA (C). The putative leader peptide was outlined in grey, along with its putative RBS sequence (blue); the *erm*(55) start codon was outlined in green along with its putative RBS (orange).

Interestingly, the 319-bp *marR*-*erm*(55) intergenic region was identical in 38/40 plasmids. The *erm*(55)^P^ downstream region up to *parA* intergenic region varied in size (1,075 to 1,332 bp) and three plasmids harboured insertion sequences: *istA* IS21 family transposase and *istB* IS21-like element helper ATPase. The best BLAST hits, excluding other plasmids, of the intergenic region upstream of *erm* were found in *M. koreensis* JCM19956 strain, *P. equi* strain PAM2287 pRErm46 plasmid, *Rhodococcus sp.* PD04 strain plasmid and *P. equi* lh_12 plasmid. The alignment of *erm*(55)^P^ intergenic regions to six mycobacterial *erm* upstream regions, including *M. koreensis* JCM19956 and *P. equi* upstream *erm*(46) regions, showed a diverse pattern of nts composition and sizes (ranges 123-295 nts, Figure 4B) depending on the species. Only one ORF of 14 codons (Met-Gly-Ile-Thr-Ser-Val-Ala-Val-Ile-Arg-Ile-Arg-Arg-Phe-Stop) was compatible with a leader peptide (*i.e.* a short, translated peptide in positive strand with a codon start not overlapping *erm*) within the *erm*(55)^P^ 169 nts upstream region (Table S3). A downstream segment was predicted to adopt a stable secondary structure (MFE = − 51.2 kcal/mol) comprising one stem loop and one double branched loop (Figure 4 C) that potentially sequester a putative RBS sequence (5’-CCCGGAGTCCG-3’). This may prevent the translation of the AUG start codon of *erm*(55)^P^. Pairwise BLAST alignments with *M. koreensis* (135/169 nts, 80%), and *P. equi* pRErm46 (125/169 nts, 74%) reveal sequence conservation of the 3′ end of this structured region, containing the putative leader peptide, stem loops, and the putative RBS (5’-CCCGGAGTGCG-3’), −10 nts from the *erm* start codon.

## Discussion

This study provides the first detection in Europe of *erm*(55)-carrying plasmids associated with macrolide inducible resistance in *M. chelonae* and other RGM species. We also provide insights regarding the species and plasmids encoding this new *erm* gene, the conserved functional domains of *erm* encoded proteins in mycobacteria, and the possible *erm*(55)^P^ regulatory region responsible for the inducible resistance.

Plasmid-borne *erm*(55)-mediated inducible macrolide resistance was first reported in the USA in 2023 and later in Japan in 2026 [4,5]. The detection of plasmid-borne *erm*(55)^P^ in France in isolates from 2016 extends the known geographic distribution, suggesting a broader distribution than previously appreciated although the temporal trend is unknown. Although direct functional validation was not performed, the concordance between *erm*(55)^P^ carriage, inducible CLR resistance, absence of *rrl* mutations, and a previous functional report [11] strongly support its role in the observed phenotype. The hypothesis of a horizontal plasmid spreading, as suggested by Graham *et al.* [6], is reinforced by the conservation of a plasmid backbone and the absence of clustering of *M. chelonae* isolates harbouring *erm*(55)^P^, although an independent acquisition from a common source cannot be excluded. *erm*(55)^P^ was reported in different RGM species, including *M. parafortuitum,* which is reported here for the first time. This suggests that even less frequently encountered RGM species may contribute to its ecological maintenance. This resistance mechanism appears to be very rare in accordance with previous studies (3.8% in the USA [6] and 0.8% in Japan [5]), and so far, only distributed in RGM. Our study was specifically targeting certain species among RGM, preventing us from determining a prevalence rate. The detection of *erm*(55)^P^ protein sequences in environmental datasets may reflect a broader ecological reservoir, but metagenomics data do not provide information on the bacterial host or genetic context of *erm*(55)^P^ detection.

Plasmids are common among RGM and SGM species [12–14]. They are inconsistently distributed, highly diverse, and have a limited number of known resistance or virulence-associated genes [13]. Closely related plasmids may be found in different species, suggesting that horizontal gene transfer may play an important role. Fortunately, *erm*(55)-carrying plasmid transfer from *M. chelonae* to *M. avium* was not identified in the patient described in our study with concomitant isolation of both species. A protein sequence showing 74% aa homology to *erm*(55)^P^ and 75% to *erm*(46) was found as a chromosomal gene in *M. koreensis,* which is an SGM species closely related to *M. triviale*, and placed between SGM and RGM in mycobacteria phylogeny [15,16]. Although the precise links between this putative *M. koreensis* new *erm* gene, *erm*(55) and *erm*(46), are not established, the *erm* integration in mycobacterial genomes is of particular concern, especially in patients with chronic NTM disease, as it creates a favourable environment for genomic exchanges.

*Bioinformatics* analysis of the plasmid sequence provided new insights into the putative mode of regulation of *erm*(55)^P^. The 23s ribosomal RNA adenine methyltransferase domain is well conserved among mycobacterial Erm proteins; however, their upstream regulatory regions vary in size and sequence. A putative 14 aa leader peptide and two consecutive hairpins mRNA secondary structure were proposed based on intergenic sequence analysis. This peptide contains consecutive Arg residues whose slow decoding by mycobacterial ribosomes in the presence of an inducer may cause ribosomal stalling and liberate the *erm*(55)^P^ RBS for productive translation. *erm*(55) 23s methyltransferase, the closest known protein is encoded by *erm*(46) from *P. equi* and other *Rhodococcus* species, which is also carried by plasmids. Regulation of *erm*(46) expression is poorly described in this genus [17], but interestingly, the 3′ −169 nts upstream *erm*(55)^P^ and *erm*(46), containing the last two stem loops, and the putative RBS shared a 74% nt identity. Although functional validation is needed to decipher *erm*(55)^P^ expression regulation, the RNA predicted secondary structure, the good conservation of the upstream intergenic region, and presence of a candidate leader peptide make it compatible with a translational attenuation mechanism. This mechanism is well described for *erm*(C) in *Staphylococcus* [18] and *erm*(B) in *Streptococcus* [19], and has been recently suggested for *Helcococcus kunzii erm*(47) [20]. In mycobacteria, *erm*-mediated macrolide resistance appears to be regulated predominantly through an inducible increase in gene expression. For *M. tuberculosis*, exposure to CLR increases *erm*(37) transcript levels, indicating inducible regulation at the RNA level [21]. Similarly, *erm(38)* mRNA level from *M. smegmatis* increases more than tenfold following macrolide exposure [22]. *erm*(38), *erm*(39), and *erm*(40) are near *folD* although the precise molecular mechanisms controlling their expression remain poorly characterised [23]. The best-understood system is *erm*(41) in *M. abscessus*, where macrolide exposure activates WhiB7 transcription factor, which induces *erm(41)* expression through a specific upstream regulatory motif [3,24].

In several mycobacterial infections, macrolides represent the only orally available agents with reliable activity, making resistance to this antibiotic class a major determinant of clinical outcome. Inducible ribosomal RNA methylation mediated by *erm* genes is an important cause of macrolide resistance, orienting the therapeutic choice [1,2]. *M. chelonae* was so far regarded as fully susceptible to macrolides, and lacking a functional *erm* gene, accordingly [3]. Although rare, *erm(55)*-mediated resistance may remain undetected if CLR MIC is interpreted only at day 3-4 [5,6], as recommended by current CLSI guidelines [25,26]. By extending the geographical distribution of *erm*(55) mediated resistance, this study aligns with a previous report to support a 14-day incubation prolongation for CLR MIC reading for *M. chelonae* and other RGM known as lacking chromosomal *erm* [6]. *erm(55)* RT-PCR may provide a useful rapid screening tool for phenotypic susceptibility testing. *erm*(55) plasmid-mediated macrolide resistance was also present in *M. neoaurum*, *M. iranicum*, *M. bacteremicum*, *M. obuense*, *M. murale* and now *M. parafortuitum*. These species, often considered as contamination or colonisation in pulmonary samples, may still be epidemiologically relevant as carriers of transferable macrolide resistance determinants.

Our study has several limitations. First, the lack of experimental expression assays prevents definitive conclusions regarding the exact regulatory mechanism of *erm*(55)^P^. The proposed attenuation-like regulatory mechanism is supported by sequence conservation and structural predictions, but further transcriptional or translational assays are needed to validate this hypothesis. Furthermore, because the screening strategy was restricted to selected species and/or isolates meeting predefined phenotypic criteria, the prevalence of *erm*(55)^P^ in the broader NTM population may have been underestimated. Additionally, limited data on treatment and outcomes for all patients hinder a comprehensive evaluation of the impact of *erm(55)^P^* carriage on therapeutic response. We could not determine the transmission route due to a lack of evidence that the patients had travelled to the US or Japan.

## Conclusions

This study reports first identification in Europe of plasmid-borne *erm*(55)^P^ in *M. chelonae* and other RGM associated with inducible macrolide resistance, and the first report in *M. parafortuitum*. Although further studies are required to clarify the *erm*(55)^P^ regulatory mechanism and the prevalence and transferability of *erm*(55)^-^carrying plasmids, our findings strengthen the need for prolonged CLR MIC reading in RGM and plasmid-mediated resistance surveillance in NTM.

## Disclosure statement

All other authors declare that they have no conflicts of interest

## Declaration of Generative AI Use

Copilot (Microsoft 365) was used for language refinement. The authors confirm the originality and accuracy of the translation, taking full responsibility for the manuscript.

## Funding

This work was supported by an annual grant from Santé Publique France (SPF) and Direction Générale de l’Offre de Soins (DGOS) for the associate laboratory of the National Reference Centre for Mycobacteria and Antimycobacterial Resistance (CNR-MyRMA). The funders had no role in study design, data collection, or the decision to submit the work for publication.

## Acknowledgments

We thank Odile Vissouarn, Célia Sicard, Nicolas Gille, Natacha Tytus and Zineb Bougadouha for technical assistance. We thank all clinical microbiologists who submitted the NTM isolates and all clinicians in charge of these patients. We thank all members of CNR-MyRMA for their contributions to this study.

Access to data

Raw reads from the five *erm*(55)^P^ carrying isolates (accession numbers PRJNA1490064, BioProject, and SAMN61416491 to SAMN61416495 BioSamples) and *erm*(55)^P^ carrying plasmids assemblies (SUB16311836, SUB16313607, SUB16313744, SUB16313779, and SUB16313804) will be released upon publication.

## Authorship contribution statement

SA & EC identified the initial findings. CA, ZA, and NB performed sequencing and MIC experiments. YC, SA, and CA conducted bioinformatics analyses. CA interpreted the bioinformatics results and wrote the original manuscript. TF and LG developed the erm(55) RT-PCR assay. CP and NV provided the M. avium genomes and participated in the CNR-MyRMA. NV was the clinician responsible for the patient with the Mc1 isolate. NB collected routine identification data. FM and EC contributed expertise on CLR resistance in RGM. EC supervised the study and reviewed the manuscript. All authors reviewed the final manuscript.

## Notes

### Competing Interest Statement

The authors have declared no competing interest.

