## Supplementary materials for "Emergence of plasmid-borne *erm*(55)-associated macrolide resistance in *Mycobacterium chelonae* and other rapidly-growing non-tuberculous mycobacteria in Europe"

**Table S1: List of *erm*(55) carrying plasmids core genes and accessory genes identified by Panaroo**

See dedicated tabular excel file

**Table S2: Comparison of the five *erm*(55) carrying plasmids found in this study with previously identified plasmids**

| ***Mycobacterium* Isolates** | | | | **Plasmid** | | | | | | | |
| --- | --- | --- | --- | --- | --- | --- | --- | --- | --- | --- | --- |
| **Species** | **Country** | **Isolate ID** | **Name** | | **Size (bp)** | **Collection Year** | **Clinical Source** | **Source Country/State: region** | **Nucleotide Accession** | **Query cover (%)** | **Homology (%)** |
| *M. chelonae* | France | Mc1 | pErm55Mc1 | | 126 019 | 2020 | Sputum | France: Paris | SUB16311836* | NA | NA |
| *M. neoaurum* | France | Mn1 | pErm55Mn1 | | 128 376 | 2023 | Blood | France: Marseille | SUB16313607* | 98 | 100 |
| *M. neoaurum* | France | Mn2 | pErm55Mn2 | | 165 340 | 2025 | Blood | France: Lille | SUB16313744* | 76 | 100 |
| *M. neoaurum* | France | Mn3 | pErm55Mn3 | | 138 589 | 2023 | Sputum | France: Paris | SUB16313779* | 91 | 100 |
| *M. parafortuitum* | France | Mp1 | pErm55Mp1 | | 127 642 | 2016 | Sputum | France: Hauts de France region | SUB16313804* | 91 | 100 |
| *M. chelonae* | USA | MCHL20831 | p20831 | | 137 526 | 2020 | Respiratory | Massachusetts | PV530475 | 92 | 99.93 |
| *M. bacteremicum* | USA | MBAC22545 | p22545 | | 156 532 | 2022 | Sputum | North Dakota | PV682378 | 81 | 99.99 |
| *M. bacteremicum* | USA | MBAC99676 | p99676 | | 156 585 | 1999 | Blood | Texas | PV682379 | 81 | 99.9 |
| *M. chelonae* | USA | MCHL18529 | p18529 | | 137 350 | 2018 | Respiratory | Washington | PV682380 | 92 | 99.92 |
| *M. chelonae* | USA | MCHL19973 | p19973 | | 138 537 | 2019 | Hand | Kentucky | PV682381 | 91 | 99.82 |
| *M. chelonae* | USA | MCHL20135 | p20135 | | 136 944 | 2020 | Toe (Pus) | Massachusetts | PV682382 | 92 | 99.98 |
| *M. chelonae* | USA | MCHL21627 | p21627 | | 136 703 | 2021 | Tissue (Knee) | Massachusetts | PV682383 | 92 | 99.51 |
| *M. chelonae* | USA | MCHL22950 | p22950 | | 163 138 | 2020 | Leg | Pennsylvania | PV682384 | 77 | 99.99 |
| *M. grossiae* | USA | MGRO13830 | p13830 | | 140 353 | 2013 | Sputum | Connecticut | PV682385 | 90 | 99.9 |
| *M. iranicum* | USA | MIRA14376 | p14376 | | 125 996 | 2014 | Blood | Ohio | PV682386 | 100 | 99.7 |
| *M. iranicum* | USA | MIRA22753 | p22753 | | 126 859 | 2022 | Leg | unknown | PV682387 | 97 | 99.99 |
| *M. neoaurum* | USA | MNEA04327 | p04327 | | 149 111 | 2004 | Sputum | New York | PV682388 | 85 | 99.98 |
| *M. neoaurum* | USA | MNEA05280 | p05280 | | 139 878 | 2005 | Blood | North Carolina | PV682389 | 90 | 99.99 |
| *M. neoaurum* | USA | MNEA06353 | p06353 | | 136 344 | 2006 | Sputum | Illinois | PV682390 | 93 | 99.99 |
| *M. neoaurum* | USA | MNEA13288 | p13288 | | 139 734 | 2013 | Blood | North Carolina | PV682391 | 90 | 99.97 |
| *M. neoaurum* | USA | MNEA14154 | p14154 | | 126 243 | 2014 | Hip | North Carolina | PV682392 | 100 | 99.89 |
| *M. neoaurum* | USA | MNEA23387 | p23387 | | 141 959 | 2023 | Blood | Kentucky | PV682393 | 89 | 99.98 |
| *M. neoaurum* | USA | MNEA23912 | p23912 | | 138 401 | 2023 | Sputum | California | PV682394 | 91 | 99.99 |
| *M. neoaurum* | USA | MNEA24983 | p24983 | | 131 018 | 2024 | Blood | Ohio | PV682395 | 96 | 99.99 |
| *M. neoaurum* | USA | MNEA97382 | p97382 | | 119 393 | 1997 | Sputum | California | PV682396 | 100 | 99.88 |
| *M. obuense* | USA | MOBU17061 | p17061 | | 140 603 | 2017 | Sputum | Massachusetts | PV682397 | 90 | 99.88 |
| *M. obuense* | USA | MOBU20363 | p20363 | | 171 667 | 2023 | Sputum | Pennsylvania | PV682398 | 70 | 99.03 |
| *M. obuense* | USA | MOBU20523 | p20523 | | 126 296 | 2023 | Eye | Iowa | PV682399 | 100 | 99.93 |
| *M. obuense* | USA | MOBU20738 | p20738 | | 137 284 | 2020 | Sputum | Minnesota | PV682400 | 92 | 99.92 |
| *M. obuense* | USA | MOBU20810 | p20810 | | 140 703 | 2020 | Wound (Foot) | Kansas | PV682401 | 90 | 99.99 |
| *M. obuense* | USA | MOBU21749 | p21749 | | 153 776 | 2021 | Finger | Minnesota | PV682402 | 82 | 99.98 |
| *M. obuense* | USA | MOBU21750 | p21750 | | 172 938 | 2021 | Sputum | Minnesota | PV682403 | 73 | 99.99 |
| *M. obuense* | USA | MOBU21751 | p21751 | | 161 529 | 2021 | Sputum | Minnesota | PV682404 | 78 | 99.87 |
| *M. obuense* | USA | MOBU22811 | p22811 | | 141 875 | 2022 | Respiratory | Washington | PV682405 | 89 | 99.98 |
| *M. iranicum* | USA | MTOR20745 | p20745 | | 122 565 | 2020 | Eye | Illinois | PV682406 | 100 | 99.98 |
| *M. chelonae* | Japan | SRL2021-127 | pErm55Mc1 | | 126 187 | 2021 | Missing | Japan:Shiga | LC872744 | 100 | 99.7 |
| *M. obuense* | Japan | SRL2019-498 | pErm55Mo1 | | 155 889 | 2019-10 | Exudate | Japan:Kanagawa, Atsugi | LC872745 | 81 | 100 |
| *M. obuense* | Japan | SRL2021-291 | pErm55Mo2 | | 170 220 | 2021 | Missing | Japan:Oita | LC872746 | 74 | 100 |
| *M. obuense* | Japan | SRL2023-024 | pErm55Mo3 | | 146 449 | 2023 | Blood | Japan:Aichi, Okazaki | LC872747 | 86 | 100 |
| *M. murale* | Japan | SRL2023-035 | pErm55Mm1 | | 143 802 | 2023-07 | Sputum | Japan:Ibaraki | LC872748 | 88 | 100 |

***** Accession number in progress

**Table S3: ORF found within *erm*(55)^P^ 319 bp upstream intergenic region using NCBI ORF finder tool**

| **Label** | **Strand** | **Frame** | **Start** | **Stop** | **Length (nt \| aa)** |
| --- | --- | --- | --- | --- | --- |
| ORF1 | + | 1 | 7 | 39 | 33 \| 10 |
| ORF2 | + | 1 | 82 | >168 | 87 \| 28 |
| ORF3 | + | 3 | 15 | 59 | 45 \| 14 |
| ORF4 | + | 3 | 84 | >167 | 84 \| 27 |
| ORF5 | - | 1 | 46 | >2 | 45 \| 14 |
| ORF6 | - | 2 | 147 | 100 | 48 \| 15 |
| ORF7 | - | 2 | 75 | 22 | 54 \| 17 |
| ORF8 | - | 3 | 149 | >3 | 147 \| 48 |

The 1^st^ position refers to the nt immediately after to *marR* stop codon and the 319 to the last nt immediately before *erm*(55). nt: nucleotide; aa: amino acid
